# Macrophage inhibitor clodronate enhances liver transduction of lentiviral but not AAV vectors or mRNA lipid nanoparticles *in vivo*

**DOI:** 10.1101/2023.07.26.550697

**Authors:** Loukia Touramanidou, Sonam Gurung, Claudiu A. Cozmescu, Dany P. Perocheau, Dale Moulding, Deborah Ridout, Alex Cavedon, Summar Siddiqui, Lisa Rice, Patrick F. Finn, Paolo G.V. Martini, Andrea Frassetto, Simon N. Waddington, John R. Counsell, Paul Gissen, Julien Baruteau

## Abstract

Recently approved adeno-associated viral (AAV) vectors for liver monogenic diseases hemophilia A and B are exemplifying the success of liver-directed viral gene therapy. In parallel, additional strategies are rapidly emerging to overcome some inherent AAV limitations, such as non-persistence of episomal transgene in rapidly growing liver and immune response. Integrating lentiviral vectors and non-viral lipid nanoparticles encapsulating mRNA (LNP-mRNA) are rapidly being developed, currently at preclinical and clinical stages respectively. Macrophages are first effector cells of the innate immune response triggered by gene therapy vectors. Macrophage uptake and activation following administration of viral gene therapy and LNPs has been reported. In this study, we assessed the biodistribution of AAV, lentiviral and LNP-mRNA gene therapy following inhibition of tissue macrophages by clodronate liposomes in neonatal and juvenile mice. Juvenile clodronate-treated mice showed significant increase of lentiviral-transduced hepatocytes, and increasing trend of transduction was shown in neonatally-injected mice. In contrast, AAV- and LNP-mRNA-treated neonatal and juvenile animals did not show significant increase of liver biodistribution following clodronate administration. These findings will have translational application for liver-targeting gene therapy programmes.

## Introduction

Over the last two decades, gene therapy has transformed the therapeutic landscape of liver monogenic diseases demonstrating maturity with numerous clinical successes [1–3]. Adeno-associated viral (AAV) vectors represent the leading gene therapy strategy for targeting liver showing a favourable outcome between safety and efficacy, especially in adult patients [4–8]. However, systemic administration of high doses of AAV vectors have shown limitations caused by severe innate and adaptive immune responses [9–12], preventing re-injections in humans [13]. AAV vectors deliver mainly episomal transgenes, which are not passed to daughter cells during cell division and liver growth [14–16]. Therefore, sustained clinical efficacy in a rapidly growing paediatric liver requires alternative gene therapy strategies such as integrative approaches, e.g. *in vivo* lentiviral vectors [17, 18], gene integration mediated by nucleases [19, 20] or non-viral technologies [21], e.g., lipid nanoparticles (LNP) encapsulating mRNA (LNP-mRNA) [22–27] respectively.

Whatever the chosen gene therapy strategy, methods to optimise hepatocyte transduction are essential for efficacy, safety and cost-effectiveness. In addition, administering a minimal effective dose improves the safety profile as some viral vectors have shown dose-dependent severity of adverse events [28]. Macrophages are the first effector cells for innate immunity. As such, liver-targeting lentiviral gene therapy *in vivo* has shown high uptake by macrophages in the splenic marginal zone reducing efficacy [17, 18, 29, 30]. Macrophage activation following AAV vector administration has been reported [31]. Additionally, LNPs can trigger innate immunity by uptake from antigen presenting cells [32].

Clodronate (dichloroethylene-bisphosphonate or Cl2MBP) is a bisphosphonate molecule with market authorisation in cancer. Clodronate-encapsulated liposomes achieve a transient depletion of circa 90% macrophages in both red pulp of the spleen and Kupffer cells in the liver at 24 hours after systemic injection (**Supplementary Figure 1**) [33, 34]. Adenoviral and AAV vectors result in the activation of the innate immune system leading to elimination of transduced cells [35–42]. Resident hepatic and splenic macrophages act as triggers of the initial non-specific immune response against pathogens and are accountable for the majority of absorbed vector particles [17, 29, 30, 43–46]. Pre-administration of clodronate liposomes followed by administration of adenoviral vectors depleted macrophages and allowed higher liver transduction *in vivo* [47–49]. In contrast, pre-administration of clodronate and AAV vector injection *in vivo* produced a considerable reduction in transgene expression in the liver [50].

Here we tested the effect of clodronate-mediated macrophage inhibition on liver transduction *in vivo* in neonatal and juvenile mice prior to administration of three different gene therapy modalities: lentiviral vector, AAV vector, and non-viral LNP-mRNA. We show that the induction of macrophage depletion through systemic administration of clodronate liposomes increases hepatocyte transduction by lentiviral vector but has no benefit for AAV vector and LNP-mRNA.

## Results

### Macrophage inhibition enhances lentiviral liver transduction in juvenile mice

Transient macrophage depletion by systemic pre-administration of clodronate increases adenoviral-mediated liver transduction [51]. We therefore assessed liver transduction after systemic administration of clodronate prior to intravenous lentiviral injection in neonatal and juvenile wild-type mice. CD1 mice received repeated intraperitoneal injections of clodronate-or PBS-encapsulated liposomes at 30 and 6 hours before a single intravenous injection with CCL.LP1.GFP vector at the dose of 4e10TU/Kg. Untreated animals were used as negative controls. One month following vector injection, mice were harvested, and livers were collected for analysis (**Figure 1A**).

**Figure 1.**
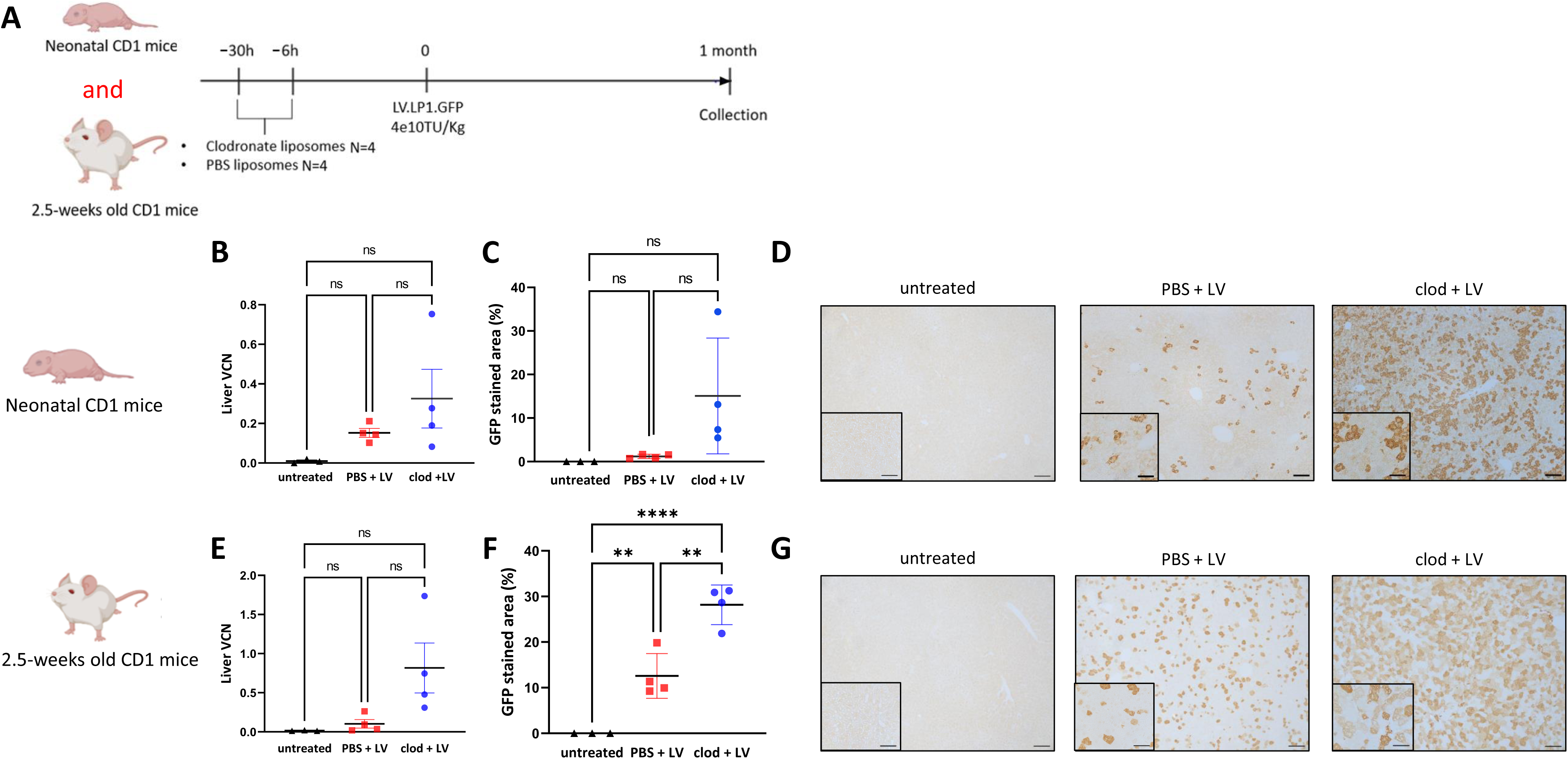
Macrophage inhibition enhances lentiviral liver transduction in juvenile mice. **(A)** Schematic representation of the experimental design testing lentiviral vector transduction following pre-treatment with clodronate liposomes in CD1 mice. **(B)** Lentiviral vector genome copies per cell in liver, **(C)** quantification of GFP immunostaining, **(D)** representative images of GFP immunostaining in liver sections of neonatally-injected CD1 mice. **(E)** Lentiviral vector genome copies per cell in liver, **(F)** quantification of GFP immunostaining, **(G)** representative images of GFP immunostaining in liver sections of 2.5-weeks-old-injected CD1 mice. **(B,C,E,F)**: Horizontal lines display the mean ± standard deviation. One-way ANOVA with Tukey’s multiple comparisons test, ns: not significant, ** *p*<0.01, **** *p*<0.0001; untreated (n=3), PBS + LV (n=4), clod + LV (n=4). **(D,G)**: Scale bars are 100μm and 50μm for x10 and x20 magnification, respectively. Clod: clodronate liposomes; LV: lentivirus; PBS: Phosphate Buffer Solution; VCN: vector copy number.

In neonates, liver vector genome copy number (VCN) showed an increasing trend of liver transduction in the clodronate-versus PBS-injected group (**Figure 1B**). Liver immunostaining also showed an increasing trend in clodronate-versus PBS-treated animals with 15% and 1.2% of GFP expression respectively (**Figure 1C**). The pattern of hepatocyte transduction revealed a homogeneous and scattered expression in all injected mice with no predominant expression in periportal or pericentral hepatocytes (**Figure 1D**). In juvenile mice, liver VCN showed an increasing trend of liver transduction in clodronate-versus PBS-group (**Figure 1E**). These results were supported by a significant increase of liver transduction of GFP immunostaining in clodronate-versus PBS-treated cohorts with 28% and 12% GFP expression respectively (*p*=0.002) (**Figure 1F**, 1G). Overall, these findings demonstrated an enhanced liver lentiviral-mediated transduction after pre-treatment with clodronate in both neonatal and juvenile animals, and a significantly increased transduction of liver in juvenile compared to neonates.

### Macrophage inhibition decreases splenic transduction and enhances lentiviral-mediated liver transduction

To assess reproducibility of enhanced liver transduction mediated by lentiviral vector following clodronate pre-exposure, the experiment performed in outbred CD1 mice was replicated with inbred C57BL/6J mice, another common mouse background strain used in research (**Figure 2A**) [52–54]. CD1 mice had been initially chosen for their superior breeding, large litters, and cost-effectiveness [55, 56]. We also assessed spleen VCN as an indirect marker of systemic macrophage depletion as previously published [17]. We confirmed in neonates the increasing trend of liver VCN (**Figure 2B**) and reducing trend of spleen VCN (**Figure 2C**) in the clodronate-versus PBS-treated group. PBS-treated group showed values similar to the untreated negative control values. GFP liver immunostaining further showed an increasing trend in the clodronate-versus PBS-treated group (**Figure 2D**, **2E**). Compared to PBS-treated group, the clodronate-treated juvenile-injected animals showed significantly increased liver VCN (*p*=0.008) (**Figure 2F**), decreased splenic VCN (*p*=0.0002) (**Figure 2G**), and increased liver GFP immunostaining (*p*=0.006) (**Figure 2H, 2I**). As observed in CD1 mice, the benefit of clodronate pre-treatment in lentiviral-mediated liver transduction was higher in juvenile compared to neonatal C57BL/6J mice. These results demonstrate that by reducing macrophage uptake of lentiviral particles via clodronate pre-treatment can enhance lentiviral mediated liver transduction. The significantly enhanced liver transduction was observed in juvenile mice in both outbred and inbred strains with comparable levels and effect observed for both liver VCN and immunostaining.

**Figure 2.**
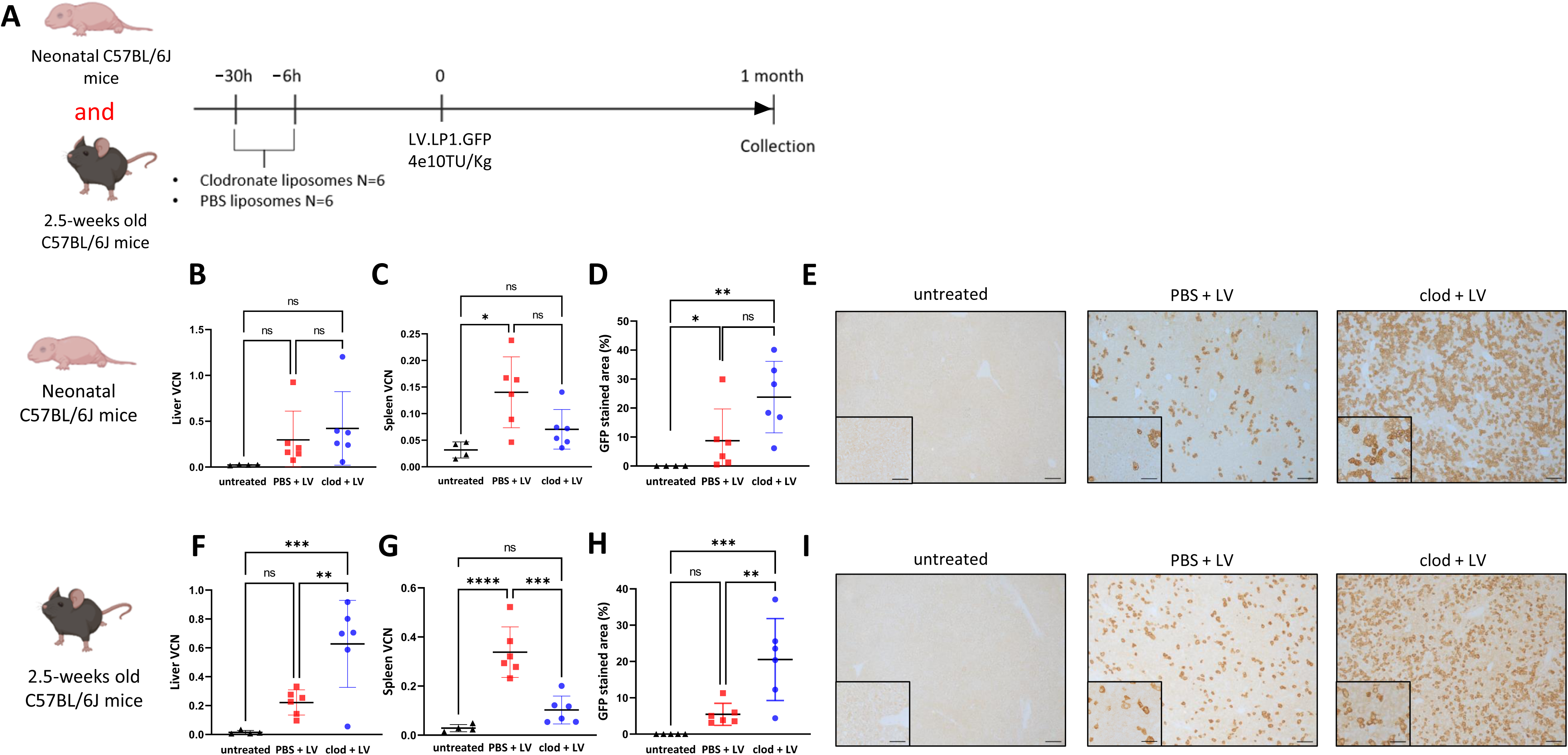
Macrophage inhibition decreases splenic transduction and enhances lentiviral-mediated liver transduction. **(A)** Schematic representation of the experimental design testing lentiviral vector transduction following pre-treatment with clodronate liposomes in CD1 mice. **(B)** Lentiviral vector genome copies per cell in liver, **(C)** vector genome copies per cell in spleen; **(D)** quantification of GFP immunostaining, **(E)** representative images of GFP immunostaining in liver sections of neonatally-injected CD1 mice. **(F)** Lentiviral vector genome copies per cell in liver, (**G**) vector genome copies per cell in spleen; **(H)** quantification of GFP immunostaining, **(I)** representative images of GFP immunostaining in liver sections of 2.5 weeks old-injected CD1 mice. **(B-H)**: Horizontal lines display the mean ± standard deviation. One-way ANOVA with Tukey’s multiple comparisons test, ns: not significant, * p<0.05, ** *p*<0.01, *** *p*<0.001, **** *p*<0.0001; untreated (n=4-5), PBS + LV (n=6), clod + LV (n=6). **(E,I)**: Scale bars are 100μm and 50μm for x10 and x20 magnification, respectively. Clod: clodronate liposomes; LV: lentivirus; PBS: Phosphate Buffer Solution; VCN: vector copy number.

### Macrophage inhibition does not influence AAV-mediated liver transduction

Neonatal and juvenile CD1 mice received intraperitoneal injections of clodronate or PBS liposomes at 30 and 6 hours before they received intravenous injection of 1e13VG/Kg of AAV8.LP1.GFP vector. Untreated animals were used as negative controls. Animals were harvested at 4 weeks post-AAV administration (**Figure 3A**).

**Figure 3.**
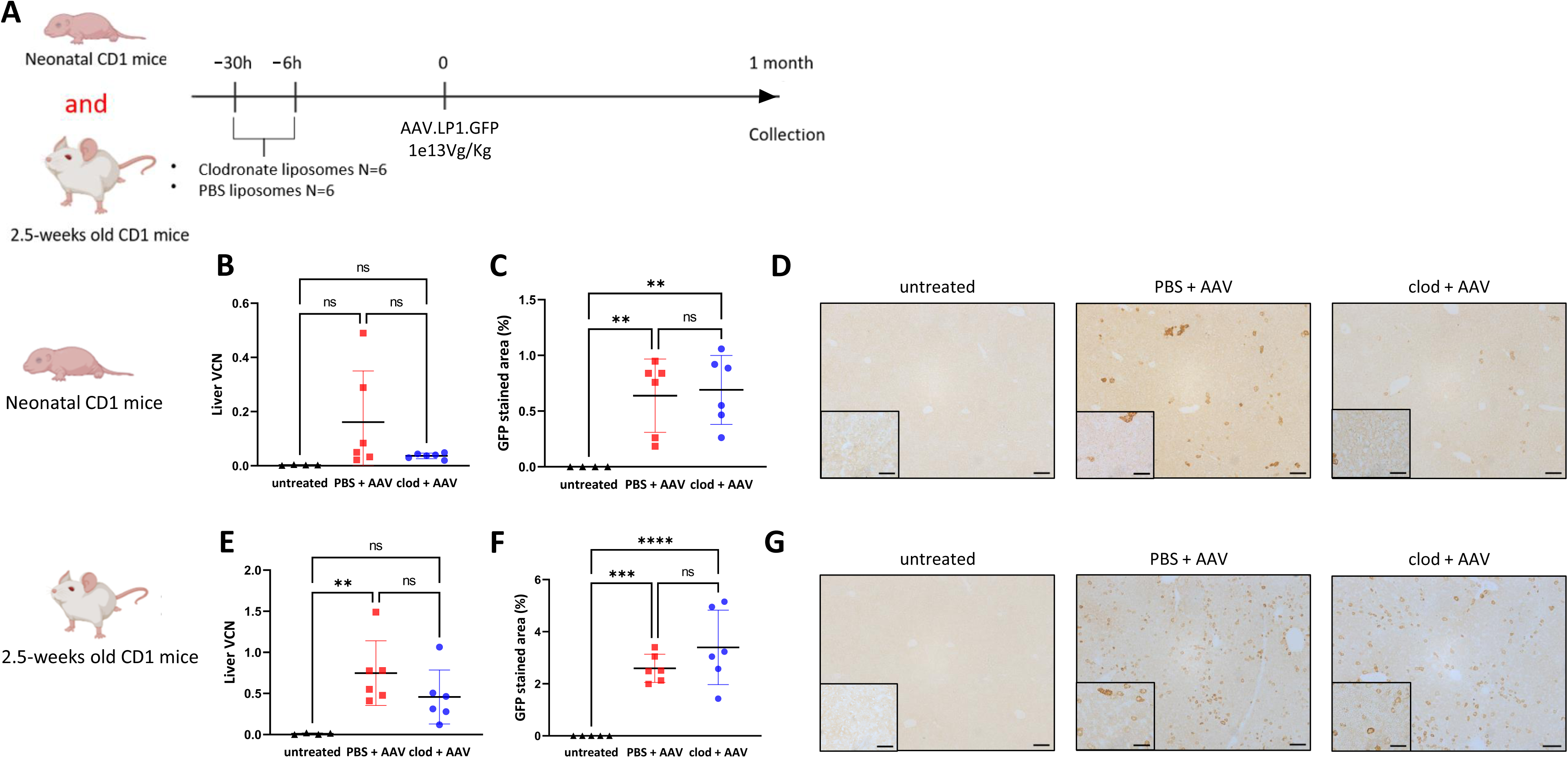
Macrophage inhibition does not influence AAV-mediated liver transduction. **(A)** Schematic representation of the experimental design testing AAV vector transduction following pre-treatment with clodronate liposomes in CD1 mice. **(B)** AAV vector genome copies per cell in liver, **(C)** quantification of GFP immunostaining, **(D)** representative images of GFP immunostaining in liver sections of neonatally-injected CD1 mice. **(E)** AAV vector genome copies per cell in liver, **(F)** quantification of GFP immunostaining, **(G)** representative images of GFP immunostaining in liver sections of 2.5 weeks old-injected CD1 mice. **(B,C,E,F)**: Horizontal lines display the mean ± standard deviation. One-way ANOVA with Tukey’s multiple comparisons test, ns: not significant, ** *p*<0.01, *** *p*<0.001, **** *p*<0.0001; untreated (n=4-5), PBS + AAV (n=6), clod + AAV (n=6). **(D,G)**: Scale bars are 100μm and 50μm for x10 and x20 magnification, respectively. AAV: adeno-associated virus, Clod: clodronate liposomes; PBS: Phosphate Buffer Solution; VCN: vector copy number.

In neonates, liver VCN and GFP immunostaining showed similar results between clodronate- and PBS-treated control groups (**Figure 3B-D**). In juvenile animals, liver VCN and GFP immunostaining did not show significant differences between clodronate-versus PBS-treated groups (**Figure 3E-G**). The GFP immunostaining was <1% and 2.4% in neonates and juvenile animals, respectively. These results are consistent with AAV-mediated episomal transgene biodistribution in rapidly growing livers, with presence of clusters of transduced hepatocytes likely associated with rare integration events. Overall, these data show no benefit of clodronate pre-treatment and transient macrophage depletion for AAV-mediated hepatocyte transduction.

### LNP-mRNA mediated liver transduction does not benefit from macrophage inhibition

Although LNPs naturally accumulate in the liver following systemic administration [26], there is still a lack of understanding of how LNPs could interact with Kupffer cells and splenic resident macrophages, which could result in off-target uptake. We tested the hypothesis that LNP-mRNA mediated liver transduction could benefit from clodronate pre-treatment. Neonatal and juvenile CD1 mice were pre-treated intraperitoneally with either PBS or clodronate encapsulated liposomes 30 and 6 hours before the intravenous administration of engineered LNP encapsulating GFP mRNA (**Figure 4A**). mRNA expression is transient and can occur as early as 30 minutes and have a peak of expression at 24 hours following systemic administration, followed by progressive decline in expression [2]. As such the animals were harvested at 24 hours following systemic injection of LNP-mRNA. Untreated mice were used as negative controls.

**Figure 4.**
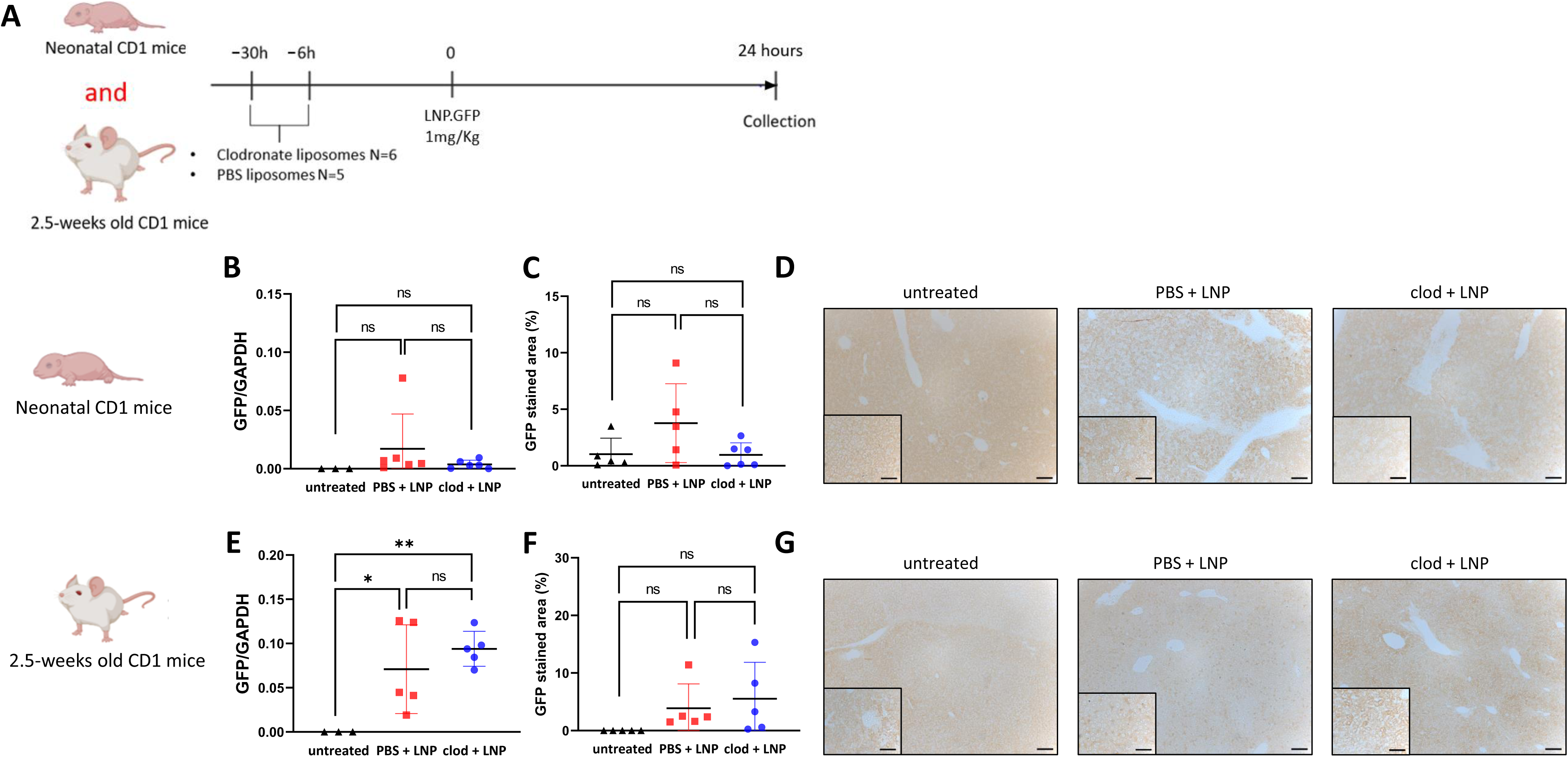
LNP-mRNA mediated liver transduction does not benefit from macrophage inhibition. **(A)** Schematic representation of the experimental design testing liver uptake of LNP.GFP following pre-treatment with clodronate liposomes in CD1 mice. **(B)** quantification of GFP western blot of livers against housekeeping control GAPDH. **(C)** quantification of GFP immunostaining, **(D)** representative images of GFP immunostaining in liver sections of neonatally-injected CD1 mice. **(E)** Quantification of GFP western blot of livers against housekeeping control GAPDH. **(F)** Quantification of GFP immunostaining, **(G)** representative images of GFP immunostaining in liver sections of juvenile-injected CD1 mice. **(B,C,E,F)**: Horizontal lines display the mean ± standard deviation. One-way ANOVA with Tukey’s multiple comparisons test, ns: not significant, *p<0.05, **p<0.01; untreated (n=3-5), PBS + LNP (n=5-6), clod + LNP (n=6). **(D,G)**: Scale bars are 100μm and 50μm for x10 and x20 magnification, respectively. Clod: clodronate liposomes; LNP: Lipid nanoparticles; PBS: Phosphate Buffer Solution.

In neonates (**Figure 4B-D**) and juvenile (**Figure 4E-G**) mice, both liver GFP western blot and immunostaining did not show enhanced expression between PBS and clodronate treated groups, suggesting no benefit of macrophage depletion in liver LNP uptake. In neonates, liver GFP immunostaining showed a decreasing trend in expression, suggesting reduced vector liver uptake. For all animals receiving LNP-mRNA, the transduction was homogeneous and diffuse. These data show that macrophage depletion has no or marginal effect on liver transduction mediated by LNP-mRNA.

## Discussion

Here we show that clodronate-mediated transient macrophage inhibition significantly increases lentiviral-mediated liver transduction in juvenile mice and shows increasing trend in neonatal mice in both outbred and inbred strains. The decreased uptake of lentiviral vectors by macrophages mechanically likely increased the vector pool for on-target liver transduction. Conversely hepatotropic AAV and LNP-mRNA did not show any enhancement in liver transduction following macrophage inhibition.

The first line of defence against viral infections consists of the innate immune response induced by the complement pathway and circulating and tissue-resident macrophages. Vesicular stomatitis virus (VSV-G)-pseudotyped lentiviral vectors are opsonised by complement-mediated inactivation in human serum, likely due to the cross-reacting, not neutralizing, and complement-fixing anti-VSV-G antibodies in humans [57–59]. Lentiviral-binding antibodies and complement proteins can opsonize lentiviral vector particles for phagocytosis by liver and spleen macrophages and professional antigen presenting cells [60–62]. Due to immune response and complement activation upon systemic administration, high amounts of lentiviral vector are uptaken by liver and splenic macrophages instead of hepatocytes., It has been shown that following intravenous injection, lentiviral vectors preferentially transduce Kupffer cells and tissue resident macrophages before hepatocytes [63]. High uptake of lentiviral vector by macrophages has previously been reported in liver and spleen, with over 70% of lentiviral DNA integrated in non-parenchymal cells in 8-week old-injected C57BL/6 mice. Fifty percent of lentiviral vector is uptaken by the spleen in non-human primates [17]. The preferential gene transfer to the spleen by VSV-G pseudotyped lentiviral vector, despite systemic administration and the well-described VSV-G pantropism, may be due to the abundant blood supply to such filtering organ [64]. Following preferential uptake by macrophages, lentiviral vectors activate the innate immune system by eliciting an inflammatory response early after vector administration [37, 65–68].

Therefore, avoiding macrophage uptake is an appealing strategy to increase the vector pool available for hepatocytes. This strategy was successfully tested *in vivo* by overexpressing the “don’t eat me” CD47 antigen signal at the capsid surface of lentiviral vectors [17]. In this study, transient depletion of macrophages by a macrophage inhibitor, clodronate liposomes, successfully benefited liver transduction mediated by lentiviral vector. Clodronate is a hydrophilic molecule that can be entrapped between concentric phospholipid bilayers to form artificial spheres or liposomes [69]. Free clodronate has a short half-life and is rapidly cleared from the circulation by the kidney while the liposome-encapsulated clodronate is preferentially uptaken by macrophages [33]. Following degradation of phospholipid bilayers by lysosomal phospholipases, clodronate is metabolised intracellularly to cytotoxic adenosine triphosphate (ATP) analogue, β,γ-Dichloromethylene ATP, leading to macrophage apoptosis [70]. Clodronate, a molecule from the bisphosphonate family, is routinely prescribed in clinical settings to prevent bone resorption in cancer [71–73]. Clodronate is generally well tolerated, with few adverse events such as transient gastrointestinal disturbances, transient increase in serum creatinine and parathyroid hormone levels [71]. This short-term and selective depletion of macrophages has shown benefit in adenoviral-mediated delivery with increased adenoviral-mediated liver transduction and reduced humoral immune response against the transgenic protein [74]. Similar to lentiviral vectors, adenoviral vectors activate strong innate immune response through both Toll-like receptor (TLR)-dependent and independent pathways resulting in upregulation of type I Interferons (IFNs) and inflammatory cytokines [75–77]. Adenoviral vectors activate the complement-mediated innate immune response via antibodies in individuals having pre-existing immunity [78].

At doses aimed for liver-targeting, AAV vectors induce a mild but detectable innate immune response. This however occurs at a lesser extent than the ones triggered by adenoviral and lentiviral vectors [79–83]. The innate immune response against AAV vectors is largely mediated by proinflammatory cytokines and chemokines in the transduced tissue as a result of TLR engagement, but is limited and highly transient [84, 85]. These molecules in turn promote immune cell induction and activation allowing the initiation and expansion of anti-transgene and/or anti-capsid adaptive immune cells, primarily CD8+ T cells [86–88]. In line with our findings, the absence of AAV-mediated enhanced liver transduction following clodronate-induced macrophage inhibition was previously reported in 8-week-old C57BL6/J mice [50]. The different observations between lentiviral and AAV-mediated liver transduction following macrophage inhibition by clodronate could be explained by the different innate immune responses triggered by each capsid.

Liver targeting typically LNPs have high affinity for hepatocytes facilitated by incorporation of apolipoprotein E (ApoE), which mediates rapid hepatocyte uptake via low-density lipoprotein receptor (LDLr) interaction [89]. As such macrophages likely play a limited role in LNP uptake or clearance. Though macrophage depletion could theoretically still provide an incremental benefit as varying lipid composition such as amino lipids, can facilitate different cell tropism within the liver microenvironment [90]. Modifying the cholesterol structure can also increase delivery to the hepatic endothelial and Kupffer cells at doses as low as 0.05mg/kg [91]. However we did not observe any enhanced liver transduction following transient macrophage inhibition, suggesting limited role of macrophage depletion on LNP mediated transduction. This could also be an indication of limited endosomal escape of the cargo which is independent of macrophage function.

In conclusion, our study shows that clodronate-induced macrophage inhibition enhances lentiviral-mediated liver transduction *in vivo*. Macrophage inhibition has no effect on AAV or LNP-mRNA mediated liver targeting gene therapy. These findings have direct translational benefit for *in vivo* lentiviral gene therapy to achieve a minimal effective dose and improve safety.

## Materials and Methods

### Experimental design

Neonatal and 2.5-week-old animals received systemic administration of clodronate liposomes (F70101C-N-FOR and F70101-NL-FOR, Stratech, Ely, UK) by repeated intraperitoneal injections at 6 and 30 hours as per manufacturer’s dose recommendation (0.2 mL for 20g animal body weight) prior to intravenous injection of viral or non-viral gene therapy vectors. Vector administration was carried out by intravenous superficial temporal vein or tail vein injection for neonatal and 2.5-weeks-old mice, respectively. Lentiviral vector dose was 4e10TU/Kg for all treated animals while 1e13Vg/Kg and 1mg/Kg was the dose for AAV and LNP-mRNA injections, respectively. Lentiviral vector was produced in house following third-generation lentiviral vector production system. Serotype 8 AAV vector was purchased by Vector Biosystems Inc (Malvern, PA, US) and LNP-mRNA were provided by Moderna Therapeutics (Massachusetts, US). Lentiviral and AAV vector-injected mice were harvested at 4 weeks following vector injection while LNP-mRNA-injected animals were harvested at 24 hours post vector injection.

### Animals

Animal procedures were approved by institutional ethical review and performed per UK home office licenses PP9223137, compliant with ARRIVE and NC3R guidelines. Wild-type C57BL/6 and CD1 mice were purchased by Charles River (Harlow, UK) and maintained on standard rodent chow (Harlan 2018, Teklab Diets, Madison, WI) with free access to water in a 12-hour light/12-hour dark environment.

### Vector production and formulation

VSV.G-pseudotyped third-generation self-inactivating (SIN) lentiviral vectors carrying the Green fluorescent protein (GFP) transgene were produced by transient transfection into HEK293T cells. Producer cells were transfected with a solution containing the selected lentiviral vector transgene backbone, the packaging plasmids pMDLg/pRRE and pCMV.REV, pMD2.G (Plasmid Factory, Bielefeld, Germany) and polyethylenimine (PEI) (24765, Polysciences, Warrington, US). Transfection media was changes after 4 hours, and supernatant was collected 48 hours after media change. Lentiviral vector-enriched supernatant was then sterilized through a 0.22μm filter and ultracentrifuged at 23,000rpm for 2 hours. Pellet containing the vector particles was then resuspended in small volumes of phosphate buffer saline (PBS), aliquoted, and stored in -80°C. After virus collection, a titration step was performed by transduction of HEK293T cells with the lentiviral vector at different dilutions. Seven days later, the transduced cells were collected, and qPCR was performed for quantification of vector genomes per mL. The AAV vector, presented the following sequence: a GFP transgene under the transcriptional activity of the LP1 promoter and with the Woodchuck Post-Regulatory Element (WPRE) downstream the transgene. GFP encoding LNP-mRNA provided by Moderna Therapeutics (Cambridge, USA) were produced using their proprietary technology.

### Vector copy number

Following liver perfusion, liver and spleen samples from lentiviral and AAV vector-injected mice were rapidly frozen using dry ice and stored at -80°C until genomic DNA extraction. The QIAgen DNeasy Blood & Tissue Kit (69504, QIAgen, Hilden, Germany) was used for genomic DNA extraction, following manufacturer’s guidelines. The plasmid standard curve was prepared by the serial dilutions ranging from 10^7^ copies/5μL to 10^3^ copies/5μL of a plasmid, containing for titin, and WPRE sequences. For lentiviral vector genome copies, the WPRE sequence was used with the following set of primers 5’-TGGATTCTGCGCGGGA -3’ (forward), 5’-GAAGGAAGGTCCGCTGGATT -3’ (reverse), 5’-FAMCTTCTGCTACGTCCCTTCGGCCCT-TAMRA -3’ (probe). The housekeeping gene *titin* was used for quantification of cell was used to normalize the results, using the following primers; for titin: 5’-AAAACGAGCAGTGACGTGAGC -3’ (forward), 5’-TTCAGTCATGCTGCTAGCGC -3’ (reverse), 5’-56-FAM/TGCACGGAAGCGTCTCGTCTCAGTC/3BHQ_1 -3’ (probe). TaqMan Universal PCR Master Mix (4304437, Thermo Fisher, Dartford, UK) was used to amplify the region of interest. The standard cycling conditions were used, starting with an initial step at 50°C for 2 minutes, followed by a 10-minute activation step at 95°C, and then 40 cycles of denaturation at 95°C for 15 seconds, annealing primers at 72°C for 1 minute, and extension at 60°C for 1 minute in a qPCR machine (4376357, Thermo Fisher, Dartford, UK).

### Immunohistochemical staining

At harvest, liver and spleen samples were fixed in 10% formalin solution, left at room temperature for 48 hours before transfer and storage in 70% ethanol at 4°C. The liver was paraffin-embedded and sectioned at 5µM thickness. The resulting slides were then kept at room temperature until staining. Sections were dewaxed in Histoclear (NAT1330, Scientific Laboratory Supplies, Nottingham, UK), dehydrated through a series graded ethanol solution to water followed by incubated in 1% H_2_O_2_ diluted in Methanol for 30 minutes to remove blood stains. Antigen retrieval was performed in boiling 0.01M citrate buffer for 20 minutes and then cooled to room temperature. Slides were blocked for non-specific binding by adding 15% goat serum (ab7481-10ml, Abcam, Cambridge, UK) diluted in 1x Tris-buffered saline with 0.1% tween-20 (TBS-T) followed by incubation in a moist chamber for 30 minutes. After washing, primary rabbit polyclonal anti-GFP (Abcam, Cambridge, UK ab290 1:1000), diluted in 10% goat serum, was added to sections, and incubated overnight at 4°C. Following 3x washing with TBS-T, 3,3’-Diaminobenzidine (DAB) staining was performed using Polink-2 Plus HRP Polymer and AP Polymer detection for Rb antibody kit (D39-18, Origene, Washington, USA) following manufacturer’s instructions. The slides were then dehydrated with increasing gradient of ethanol to water followed by a final step with Histoclear. The slides were mounted with water-free mounting medium (100579, Merk, Darmstadt, Germany) and dried overnight. Images of liver samples with DAB staining were obtained using a microscope camera (DFC420, Leica Microsystems, Milton Keynes, UK) and software (Image Analysis; Leica Microsystems, Wetzlar, Germany) was utilized to capture representative images. Quantitative analysis was performed by threshold analysis using the Image J software (Maryland, USA) (**Supplementary macro 1**).

### Western blot

30mg of liver was homogenised in ice-cold 1x RIPA buffer (Cell Signalling, Leiden, Netherlands) using Precellys homogenising tube and homogeniser, centrifuged at 10,000g for 20 minutes at 4°C. Protein concentration was measured using BCA Protein Assay kit (23227, Thermo Fisher Scientific, Dartford, UK). For each sample, 40µg of protein was diluted 1:1 with 2x Laemmli sample buffer (containing 10% 2-β-mercaptoethanol (β-ME)) making up 40µL total volume, followed by vortexing and heating to 95°C for 10 minutes. SDS-PAGE was used to separate the proteins at 100V for 1 hour and wet transfer of proteins into an immobilin PVDF membrane was performed at 400mA for 1 hour. The membrane was blocked in 5% non-fat milk powder in PBS-T followed by overnight incubation at 4°C with primary antibodies (anti-GFP; Abcam ab290 1:1000, anti-GAPDH; Abcam ab9485 1:10000, Cambridge, UK) 3x 5 minute washes with PBS-T, 1 hour incubation with fluorescent secondary antibodies (IRDye® 800CW Goat anti-Rabbit IgG 1:1000, 926-32210 and IRDye® 680RD Donkey anti-Mouse IgG, 923-68072, Licor, Nebraska, USA) and 3x 5 minutes washes with PBS-T. Image acquisition and analysis was performed using Licor Odyssey and image analysed using Licor ImageStudio Lite software (Nebraska, USA).

### Statistical analysis

Data was analysed and represented using Graphpad Prism 9.0 software (San Diego, CA, USA). Graphs display the mean ± standard deviation. Comparison were made between independent groups using one-way ANOVA with Tukey’s multiple comparisons test.

## Author Contributions

JB and LT designed the study. LT and SG conducted most of the experimental work. DP and CC, SW, DM, and DR contributed to technical assistance in experimental work. PFF, AC, SS, LR, PGVM, AF provided the *GFP* mRNA construct. LT, SG and JB wrote the manuscript. All authors reviewed and approved the manuscript.

## Acknowledgements

The authors thank Samantha Richards, Rebecca Towns, Katherine Howett, Mirabela Bandol and the staff from UCL Biological Services for their help with animal breeding and maintenance at University College London.

## References

1. Duff, C., I.E. Alexander, and J. Baruteau, Gene therapy for urea cycle defects: An update from historical perspectives to future prospects. J Inherit Metab Dis, 2023.

2. Gurung, S., et al., *mRNA therapy restores ureagenesis and corrects glutathione metabolism in argininosuccinic aciduria*. bioRxiv, 2022: p. 2022.10.19.512931.

3. Ashley, S.N., et al., Adeno-associated viral gene therapy corrects a mouse model of argininosuccinic aciduria. Molecular Genetics and Metabolism, 2018. 125(3): p. 241–250.

4. Manno, C.S., et al., Successful transduction of liver in hemophilia by AAV-Factor IX and limitations imposed by the host immune response. Nature medicine, 2006. 12(3): p. 342–347.

5. Nathwani, A.C., et al., Adenovirus-associated virus vector–mediated gene transfer in hemophilia B. New England Journal of Medicine, 2011. 365(25): p. 2357–2365.

6. George, L.A., et al., Hemophilia B gene therapy with a high-specific-activity factor IX variant. New England Journal of Medicine, 2017. 377(23): p. 2215–2227.

7. Miesbach, W., et al., Gene therapy with adeno-associated virus vector 5–human factor IX in adults with hemophilia B. Blood, The Journal of the American Society of Hematology, 2018. 131(9): p. 1022–1031.

8. Rangarajan, S., et al., AAV5–factor VIII gene transfer in severe hemophilia A. New England Journal of Medicine, 2017. 377(26): p. 2519–2530.

9. Calcedo, R., et al., Worldwide Epidemiology of Neutralizing Antibodies to Adeno-Associated Viruses. The Journal of Infectious Diseases, 2009. 199(3): p. 381–390.

10. Wang, L., et al., The Pleiotropic Effects of Natural AAV Infections on Liver-directed Gene Transfer in Macaques. Molecular Therapy, 2010. 18(1): p. 126–134.

11. Perocheau, D.P., et al., Age-Related Seroprevalence of Antibodies Against AAV-LK03 in a UK Population Cohort. Human Gene Therapy, 2018. 30(1): p. 79–87.

12. Fitzpatrick, Z., et al., Influence of Pre-existing Anti-capsid Neutralizing and Binding Antibodies on AAV Vector Transduction. Molecular Therapy – Methods & Clinical Development, 2018. 9: p. 119–129.

13. Mingozzi, F. and K.A. High, Overcoming the host immune response to adeno-associated virus gene delivery vectors: the race between clearance, tolerance, neutralization, and escape. Annual review of virology, 2017. 4: p. 511–534.

14. Penaud-Budloo, M., et al., Adeno-associated virus vector genomes persist as episomal chromatin in primate muscle. J Virol, 2008. 82(16): p. 7875–85.

15. Deyle, D.R. and D.W. Russell, Adeno-associated virus vector integration. Curr Opin Mol Ther, 2009. 11(4): p. 442–7.

16. Dalwadi, D.A., et al., AAV integration in human hepatocytes. Mol Ther, 2021. 29(10): p. 2898–2909.

17. Milani, M., et al., Phagocytosis-shielded lentiviral vectors improve liver gene therapy in nonhuman primates. Science Translational Medicine, 2019. 11(493): p. eaav7325.

18. Milani, M., et al., Liver-directed lentiviral gene therapy corrects hemophilia A mice and achieves normal-range factor VIII activity in non-human primates. Nat Commun, 2022. 13(1): p. 2454.

19. Cunningham, S.C., et al., Modeling correction of severe urea cycle defects in the growing murine liver using a hybrid recombinant adeno-associated virus/piggyBac transposase gene delivery system. Hepatology, 2015. 62(2): p. 417–28.

20. Yang, Y., et al., A dual AAV system enables the Cas9-mediated correction of a metabolic liver disease in newborn mice. Nature Biotechnology, 2016. 34(3): p. 334–338.

21. Barzel, A., et al., Promoterless gene targeting without nucleases ameliorates haemophilia B in mice. Nature, 2015. 517(7534): p. 360–4.

22. Hackett, P.B., Integrating DNA vectors for gene therapy. Mol Ther, 2007. 15(1): p. 10–2.

23. Schagen, F.H.E., et al., Insertion vectors for gene therapy. Gene Therapy, 2000. 7(4): p. 271–272.

24. Buchschacher, G.L., Jr. and F. Wong-Staal, Development of lentiviral vectors for gene therapy for human diseases. Blood, 2000. 95(8): p. 2499–2504.

25. Martini, P.G.V. and L.T. Guey, A New Era for Rare Genetic Diseases: Messenger RNA Therapy. Hum Gene Ther, 2019. 30(10): p. 1180–1189.

26. Hajj, K.A. and K.A. Whitehead, Tools for translation: non-viral materials for therapeutic mRNA delivery. Nature Reviews Materials, 2017. 2(10): p. 17056.

27. Jung, H.N., et al., Lipid nanoparticles for delivery of RNA therapeutics: Current status and the role of in vivo imaging. Theranostics, 2022. 12(17): p. 7509–7531.

28. Kishimoto, T.K. and R.J. Samulski, Addressing high dose AAV toxicity – ’one and done’ or ’slower and lower’? Expert Opin Biol Ther, 2022. 22(9): p. 1067–1071.

29. Noelia, A. and A. Castrillo, Origin and specialization of splenic macrophages. Cellular immunology, 2018. 330: p. 151–158.

30. Kashimura, M., The human spleen as the center of the blood defense system. International Journal of Hematology, 2020. 112(2): p. 147–158.

31. Muhuri, M., et al., Overcoming innate immune barriers that impede AAV gene therapy vectors. J Clin Invest, 2021. 131(1).

32. Connors, J., et al., Lipid nanoparticles (LNP) induce activation and maturation of antigen presenting cells in young and aged individuals. Communications Biology, 2023. 6(1): p. 188.

33. van Rooijen, N. and E. van Kesteren-Hendrikx, CLODRONATE LIPOSOMES: PERSPECTIVES IN RESEARCH AND THERAPEUTICS. Journal of Liposome Research, 2002. 12(1-2): p. 81–94.

34. Van Rooijen, N. and A. Sanders, Liposome mediated depletion of macrophages: mechanism of action, preparation of liposomes and applications. J Immunol Methods, 1994. 174(1-2): p. 83–93.

35. Yang, Y., et al., Cellular immunity to viral antigens limits E1-deleted adenoviruses for gene therapy. Proceedings of the National Academy of Sciences, 1994. 91(10): p. 4407–4411.

36. Yang, Y., et al., Immune responses to viral antigens versus transgene product in the elimination of recombinant adenovirus-infected hepatocytes in vivo. Gene therapy, 1996. 3(2): p. 137–144.

37. Crystal, R.G., et al., Administration of an adenovirus containing the human CFTR cDNA to the respiratory tract of individuals with cystic fibrosis. Nature genetics, 1994. 8(1): p. 42–51.

38. Zhang, H.-G., et al., Inhibition of tumor necrosis factor α decreases inflammation and prolongs adenovirus gene expression in lung and liver. Human gene therapy, 1998. 9(13): p. 1875–1884.

39. Raper, S.E., et al., A pilot study of in vivo liver-directed gene transfer with an adenoviral vector in partial ornithine transcarbamylase deficiency. Human gene therapy, 2002. 13(1): p. 163–175.

40. Tuohy, G.P. and R. Megaw, A systematic review and meta-analyses of interventional clinical trial studies for gene therapies for the inherited retinal degenerations (IRDs). Biomolecules, 2021. 11(5): p. 760.

41. Halbert, C.L., et al., Capsid-expressing DNA in AAV vectors and its elimination by use of an oversize capsid gene for vector production. Gene therapy, 2011. 18(4): p. 411–417.

42. Russell, S., et al., Efficacy and safety of voretigene neparvovec (AAV2-hRPE65v2) in patients with RPE65-mediated inherited retinal dystrophy: a randomised, controlled, open-label, phase 3 trial. The Lancet, 2017. 390(10097): p. 849–860.

43. Den Haan, J.M. and G. Kraal, Innate immune functions of macrophage subpopulations in the spleen. Journal of innate immunity, 2012. 4(5-6): p. 437–445.

44. Di Paolo, N.C., et al., Virus binding to a plasma membrane receptor triggers interleukin-1α-mediated proinflammatory macrophage response in vivo. Immunity, 2009. 31(1): p. 110–121.

45. Worgall, S., et al., Innate immune mechanisms dominate elimination of adenoviral vectors following in vivo administration. Human gene therapy, 1997. 8(1): p. 37–44.

46. Alemany, R., K. Suzuki, and D.T. Curiel, Blood clearance rates of adenovirus type 5 in mice. Journal of General Virology, 2000. 81(11): p. 2605–2609.

47. Wang, S., et al., Effect of clodronate on macrophage depletion and adenoviral-mediated transgene expression in salivary glands. J Oral Pathol Med, 1999. 28(4): p. 145–51.

48. Alzuguren, P., et al., Transient depletion of specific immune cell populations to improve adenovirus-mediated transgene expression in the liver. Liver International, 2015. 35(4): p. 1274–1289.

49. Wolff, G., et al., Enhancement of in vivo adenovirus-mediated gene transfer and expression by prior depletion of tissue macrophages in the target organ. Journal of virology, 1997. 71(1): p. 624–629.

50. Yu, D.L., et al., Macrophage Depletion via Clodronate Pretreatment Reduces Transgene Expression from AAV Vectors In Vivo. Viruses, 2021. 13(10).

51. Schiedner, G., et al., Selective depletion or blockade of Kupffer cells leads to enhanced and prolonged hepatic transgene expression using high-capacity adenoviral vectors. Molecular Therapy, 2003. 7(1): p. 35–43.

52. 52. Silver, L., Inbred Strain, in Brenner’s Encyclopedia of Genetics (Second Edition), S. Maloy and K. Hughes, Editors. 2001, Academic Press: San Diego. p. 53.

53. Casellas, J., Inbred mouse strains and genetic stability: a review. Animal, 2011. 5(1): p. 1–7.

54. 54. Roderick, T.H., Mouse, in Brenner’s Encyclopedia of Genetics (Second Edition), S. Maloy and K. Hughes, Editors. 2013, Academic Press: San Diego. p. 482–485.

55. Hsieh, L.S., et al., Outbred CD1 mice are as suitable as inbred C57BL/6J mice in performing social tasks. Neurosci Lett, 2017. 637: p. 142–147.

56. 56. Gad, S.C., Diesel Fuel, in Encyclopedia of Toxicology (Third Edition), P. Wexler, Editor. 2014, Academic Press: Oxford. p. 115–118.

57. Annoni, A., et al., Modulation of immune responses in lentiviral vector-mediated gene transfer. Cell Immunol, 2019. 342: p. 103802.

58. DePolo, N.J., et al., VSV-G pseudotyped lentiviral vector particles produced in human cells are inactivated by human serum. Mol Ther, 2000. 2(3): p. 218–22.

59. Annoni, A., et al., Liver gene therapy by lentiviral vectors reverses anti-factor IX pre-existing immunity in haemophilic mice. EMBO Mol Med, 2013. 5(11): p. 1684–97.

60. Kawai, T. and S. Akira, Innate immune recognition of viral infection. Nature immunology, 2006. 7(2): p. 131–137.

61. Brown, B.D. and D. Lillicrap, Dangerous liaisons: the role of “danger” signals in the immune response to gene therapy. Blood, The Journal of the American Society of Hematology, 2002. 100(4): p. 1133–1140.

62. Akira, S., S. Uematsu, and O. Takeuchi, Pathogen recognition and innate immunity. Cell, 2006. 124(4): p. 783–801.

63. Follenzi, A., L. Santambrogio, and A. Annoni, Immune responses to lentiviral vectors. Curr Gene Ther, 2007. 7(5): p. 306–15.

64. Finkelshtein, D., et al., LDL receptor and its family members serve as the cellular receptors for vesicular stomatitis virus. Proceedings of the National Academy of Sciences, 2013. 110(18): p. 7306–7311.

65. Benihoud, K., et al., Efficient, repeated adenovirus-mediated gene transfer in mice lacking both tumor necrosis factor alpha and lymphotoxin α. Journal of virology, 1998. 72(12): p. 9514–9525.

66. Zhang, H.G., et al., Inhibition of tumor necrosis factor alpha decreases inflammation and prolongs adenovirus gene expression in lung and liver. Hum Gene Ther, 1998. 9(13): p. 1875–84.

67. Muruve, D.A., et al., Adenoviral gene therapy leads to rapid induction of multiple chemokines and acute neutrophil-dependent hepatic injury in vivo. Human gene therapy, 1999. 10(6): p. 965–976.

68. Schnell, M.A., et al., Activation of innate immunity in nonhuman primates following intraportal administration of adenoviral vectors. Molecular Therapy, 2001. 3(5): p. 708–722.

69. Akbarzadeh, A., et al., Liposome: classification, preparation, and applications. Nanoscale Res Lett, 2013. 8(1): p. 102.

70. van Rooijen, N., A. Sanders, and T.K. van den Berg, Apoptosis of macrophages induced by liposome-mediated intracellular delivery of clodronate and propamidine. J Immunol Methods, 1996. 193(1): p. 93–9.

71. Hurst, M. and S. Noble, Clodronate. Drugs & Aging, 1999. 15(2): p. 143–167.

72. Green, J.R., Antitumor effects of bisphosphonates. Cancer, 2003. 97(3 Suppl): p. 840–7.

73. Santini, D., et al., The antineoplastic role of bisphosphonates: from basic research to clinical evidence. Ann Oncol, 2003. 14(10): p. 1468–76.

74. Schiedner, G., et al., Selective depletion or blockade of Kupffer cells leads to enhanced and prolonged hepatic transgene expression using high-capacity adenoviral vectors. Mol Ther, 2003. 7(1): p. 35–43.

75. Yamaguchi, T., et al., Role of MyD88 and TLR9 in the innate immune response elicited by serotype 5 adenoviral vectors. Human gene therapy, 2007. 18(8): p. 753–762.

76. Huang, X. and Y. Yang, Innate immune recognition of viruses and viral vectors. Human gene therapy, 2009. 20(4): p. 293–301.

77. Minari, J., S. Mochizuki, and K. Sakurai, Enhanced cytokine secretion owing to multiple CpG side chains of DNA duplex. Oligonucleotides, 2008. 18(4): p. 337–344.

78. Appledorn, D., et al., Complex interactions with several arms of the complement system dictate innate and humoral immunity to adenoviral vectors. Gene therapy, 2008. 15(24): p. 1606–1617.

79. Carestia, A., et al., Modulation of the liver immune microenvironment by the adeno-associated virus serotype 8 gene therapy vector. Mol Ther Methods Clin Dev, 2021. 20: p. 95–108.

80. Büeler, H., Adeno-associated viral vectors for gene transfer and gene therapy. Biological chemistry, 1999. 380(6): p. 613–622.

81. Kay, M.A., J.C. Glorioso, and L. Naldini, Viral vectors for gene therapy: the art of turning infectious agents into vehicles of therapeutics. Nature medicine, 2001. 7(1): p. 33–40.

82. Carter, P. and R. Samulski, Adeno-associated viral vectors as gene delivery vehicles. International journal of molecular medicine, 2000. 6(1): p. 17–44.

83. Zaiss, A.K., et al., Differential activation of innate immune responses by adenovirus and adeno-associated virus vectors. J Virol, 2002. 76(9): p. 4580–90.

84. Zhu, J., X. Huang, and Y. Yang, The TLR9-MyD88 pathway is critical for adaptive immune responses to adeno-associated virus gene therapy vectors in mice. The Journal of clinical investigation, 2009. 119(8): p. 2388–2398.

85. Zaiss, A.K., et al., Complement is an essential component of the immune response to adeno-associated virus vectors. Journal of virology, 2008. 82(6): p. 2727–2740.

86. Ertl, H.C. and K.A. High, Impact of AAV capsid-specific T-cell responses on design and outcome of clinical gene transfer trials with recombinant adeno-associated viral vectors: an evolving controversy. Human gene therapy, 2017. 28(4): p. 328–337.

87. Rogers, G.L., et al., Innate immune responses to AAV vectors. Frontiers in microbiology, 2011. 2: p. 194.

88. Hösel, M., et al., Toll-like receptor 2–mediated innate immune response in human nonparenchymal liver cells toward adeno-associated viral vectors. Hepatology, 2012. 55(1): p. 287–297.

89. Akinc, A., et al., Targeted delivery of RNAi therapeutics with endogenous and exogenous ligand-based mechanisms. Mol Ther, 2010. 18(7): p. 1357–64.

90. Sago, C.D., et al., Cell subtypes within the liver microenvironment differentially interact with lipid nanoparticles. Cellular and Molecular Bioengineering, 2019. 12: p. 389–397.

91. Paunovska, K., et al., Nanoparticles containing oxidized cholesterol deliver mRNA to the liver microenvironment at clinically relevant doses. Advanced materials, 2019. 31(14): p. 1807748.

