## Supplementary material for "Macrophage inhibitor clodronate enhances liver transduction of lentiviral but not AAV vectors or mRNA lipid nanoparticles *in vivo*"

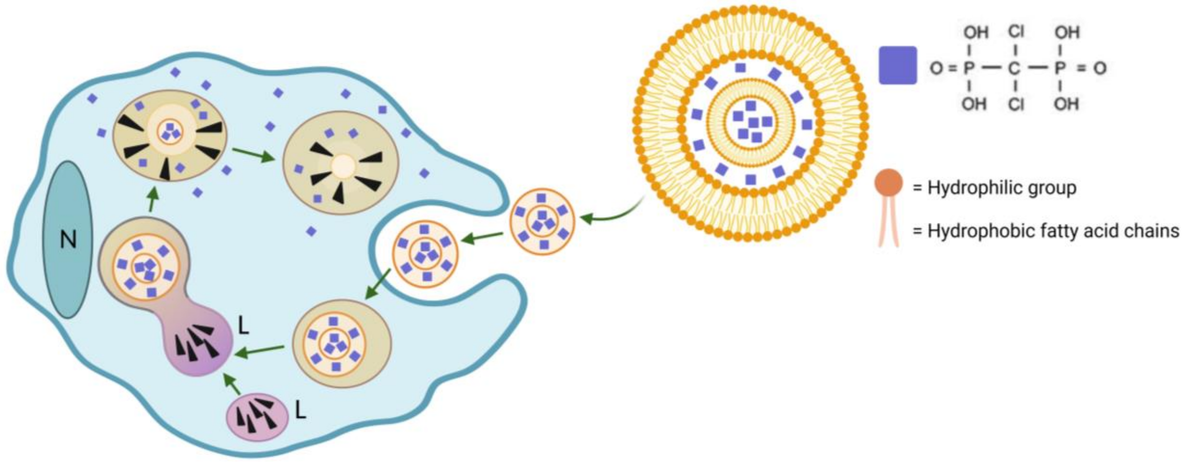


**Supplementary Figure 1. Schematic representation of clodronate liposome and its mechanism of action after macrophage uptake.** The liposome is made of concentric phospholipid bilayers, divided by aqueous compartments. The bilayers include a hydrophilic and hydrophobic group. Clodronate molecules (purple squares) are dissolved in aqueous compartments and encapsulated within the liposome. Clodronate liposomes are uptaken by macrophages via endocytosis. Late endosomes then fuse with lysosomes (L). Lysosomal phospholipases (arrowheads) digest the liposome. Clodronate will then accumulate and generate a dose-dependent toxicity ultimately triggering apoptosis. N: nucleus.

```
// macro to use color deconvolution to quantify Dab % coverage.
dir1 = getDirectory("Input folder"); //select an input folder
dir2 = getDirectory("Choose a folder to save to"); //select an output folder.
list = getFileList(dir1); //make a list of the filenames
setBatchMode(true);             //turn on batch mode so it runs in the background
// repeats the macro for every file in the folder using a for loop.
        for (i=0; i<list.length; i++) {
```

```
		showProgress(i+1, list.length);
		filename = dir1 + list[i];
				open(filename);
		name = File.nameWithoutExtension;
		rename("InputImage");
		run("Colour Deconvolution", "vectors=[H DAB] hide");
		selectWindow("InputImage-(Colour_3)");
		close();
		selectWindow("InputImage-(Colour_1)");
		close();
		selectWindow("InputImage-(Colour_2)");
		run("Median...", "radius=2"); // perhaps change this?= Gaussian?
		setThreshold(0, 140); //change this value 140. Lower value = detects more DAB.
		setOption("BlackBackground", true);
		run("Convert to Mask");
		rename(name+"-DabThreshold");
		run("Set Measurements...", "area area_fraction display redirect=None decimal=3");
		run("Analyze Particles...", "  show=Nothing summarize");
		run("Blue");  // make the overlay a colour (Red, Green, Magenta, Cyan etc)
		//run("Divide...", "value=4") // Make the overlay less bright (here by a factor of 4)
		rename("Overlay");
		run("Duplicate...", "title=outline");
		run("Outline");
		run("Red");
		selectWindow("InputImage");
```

```
			run("Add Image...", "image=Overlay x=0 y=0 opacity=25 zero"); // you can change the opacity
			selectWindow("InputImage");
			run("Add Image...", "image=outline x=0 y=0 opacity=100 zero");
			selectWindow("InputImage");
			run("Flatten");;
			saveAs("Tif", dir2+name+"-DabOutlines");//saves an image of the result
			run("Close All");
		}
selectWindow("Summary");
saveAs("Results", dir2+"All Results.csv");// save the summary as a .csv.
exit("Finished - DAB measured in "+i+" images"); // close the macro and display a window with number of images processed.
```

**Supplementary macro 1: Macro used for quantification of the DAB-stained liver sections.**
